# Reinterpretation of the localization of the ATP binding cassette transporter ABCG1 in insulin-secreting cells and insights regarding its trafficking and function

**DOI:** 10.1101/327155

**Authors:** Megan T. Harris, Syed Saad Hussain, Candice M. Inouye, Anna M. Castle, J. David Castle

## Abstract

The ABC transporter ABCG1 regulates intracellular cholesterol. We showed previously that ABCG1 deficiency inhibits insulin secretion by pancreatic beta cells and, based on its immunolocalization to insulin granules, proposed its essential role in forming cholesterol-enriched granule membranes. While we confirm elsewhere that ABCG1, alongside ABCA1 and oxysterol binding protein OSBP, supports insulin granule formation, the aim here is to update our localization and to provide added insight regarding ABCG1’s trafficking and sites of function. We show that stably expressed GFP-tagged ABCG1 closely mimics the distribution of endogenous ABCG1 in pancreatic INS1 cells and accumulates in the trans-Golgi network (TGN), endosomal recycling compartment (ERC) and on the cell surface but not on insulin granules, early or late endosomes. Notably, ABCG1 is short-lived, and proteasomal and lysosomal inhibitors both decrease its degradation. Following blockade of protein synthesis, GFP-ABCG1 first disappears from the ER and TGN and later from the ERC and plasma membrane. Beyond aiding granule formation, our findings raise the prospect that ABCG1 may act beyond the TGN to regulate activities involving the endocytic pathway, especially as the amount of transferrin receptor is increased in ABCG1-deficient cells. Thus, ABCG1 may function at multiple intracellular sites and the plasma membrane as a roving sensor and modulator of cholesterol distribution and membrane trafficking.

## Introduction

In eukaryotic cells, the ATP Binding Cassette (ABC) transporters ABCA1 and ABCG1 are known to promote cholesterol export from cells and have been of substantial interest due to their complementary roles alongside cholesterol uptake, biosynthesis and storage in maintaining intracellular cholesterol homeostasis [1,2]. While these transporters are broadly expressed, their levels are amplified in cells, e.g., macrophages and type-2 pneumocytes that are specialized for processing and exporting lipids including cholesterol physiologically [3–5], and a major focus in studying their actions has been on the mechanisms and pathways they use to transfer cholesterol to plasma lipoproteins (reviewed in [6]). Several studies have also highlighted the ability of ABCA1 and ABCG1 to promote cholesterol esterification and storage under conditions that preclude cholesterol export [7–9] and to regulate processes that do not obviously relate directly to either cholesterol export or storage including organization of plasma membrane lipids (ABCA1: [10,11]), proliferation of immune and hematopoietic cells (ABCG1: [12–14]), and insulin secretion (ABCs A1 and G1: [15–18]). ABCG1 has not been studied as extensively as ABCA1, which gained early and enduring attention due to the link between its deficiency and Tangier disease, where intracellullar cholesterol levels are increased and plasma levels of high density lipoprotein are dramatically decreased [19]. Even though ABCG1 is able to contribute to the export of cholesterol and related sterols when overexpressed or under conditions of increased cellular cholesterol load [3,8,20–24], its deficiency does not noticeably perturb cellular cholesterol levels or plasma lipoprotein profiles [3,4,16]. Thus, attention has turned to considering ABCG1’s possible intracellular roles [4].

Our laboratory participated in earlier studies of the inhibitory effect of ABCG1 deficiency on insulin secretion, and together with our colleagues, we developed the hypothesis that the transporter indeed functions intracellularly where it supports the formation of insulin granules [16]. As a part of this effort, we presented evidence using immunofluorescence microscopy and cell fractionation that ABCG1 was extensively localized to insulin granules. We now show, based on new data, that it is necessary to modify this diagnosed localization and to present a revised picture indicating that ABCG1 does not reside substantially in mature insulin granules but instead noticeably accumulates in the *trans*-Golgi network (TGN), the endosomal recycling compartment (ERC), and to a more limited extent in other cellular compartments including the plasma membrane. We reconcile its broader distribution with its relatively short lifetime by showing that ABCG1 is an itinerant protein without a persistent residence that accumulates at sites, that have been identified elsewhere as rate-limiting during membrane trafficking [25,26]. Importantly, the revised localization does not alter our hypothesis that ABCG1 supports the formation of insulin granules as we have shown in a separate study [27]. At the same time, transporter accumulation in the ERC and at the cell surface argues that ABCG1 may function more broadly and thereby tune lipid composition, especially cholesterol distribution, at multiple sites.

## Results

### ABCG1 does not appreciably co-fractionate with markers of insulin granules

As part of extending our earlier investigation of the role of ABCG1 in insulin secretion [16], we sought to learn more about its intracellular distribution. Initially, we examined the distribution of the transporter in rat insulinoma INS1 cells using subcellular fractionation by sucrose gradient centrifugation. We found that ABCG1 had a very broad distribution that did not align selectively with any individual organelle marker, suggesting that it might be present in multiple different organelles (Fig 1A). Even with this broad distribution, we were puzzled that relatively little ABCG1 appeared to co-migrate at high densities (fractions 10–12) with carboxypeptidase E (CPE) and the higher density shoulder of proinsulin’s distribution. Both of these markers are known to be present in insulin granules, particularly in immature granules where proinsulin is processed to mature insulin. Because our earlier studies were carried out using MIN6 mouse insulinoma cells, we performed a similar fractionation on them. Interestingly, the broad distribution of ABCG1 is more displaced toward higher densities than in INS1 cells, and it overlapped the distributions of proinsulin and CPE mainly in fractions 8–11. However, little ABCG1 co-fractionated with mature insulin granules (marked by insulin ELISA) that were located in fractions 12–14 (Fig 1B). In considering these results in relation to our earlier data, we realized that we had mistakenly compared the distribution of ABCG1 to proinsulin rather than to mature insulin by western blotting. Together, the previous and current findings have consistently shown that ABCG1 exhibits appreciable overlap with both proinsulin and the lower-density portions of CPE, as in Fig 1A, B, consistent with the presence of ABCG1 in the TGN and forming insulin granules but negligible overlap with fractions containing mature granules. Notably, the possible presence of ABCG1 in immature granules, particularly in MIN6 cells, is consistent with our earlier observation of a portion of ABCG1 in lower-density granules purified from islet p cells using a different fractionation procedure (Fig 4D in [16]).

**Fig 1.**
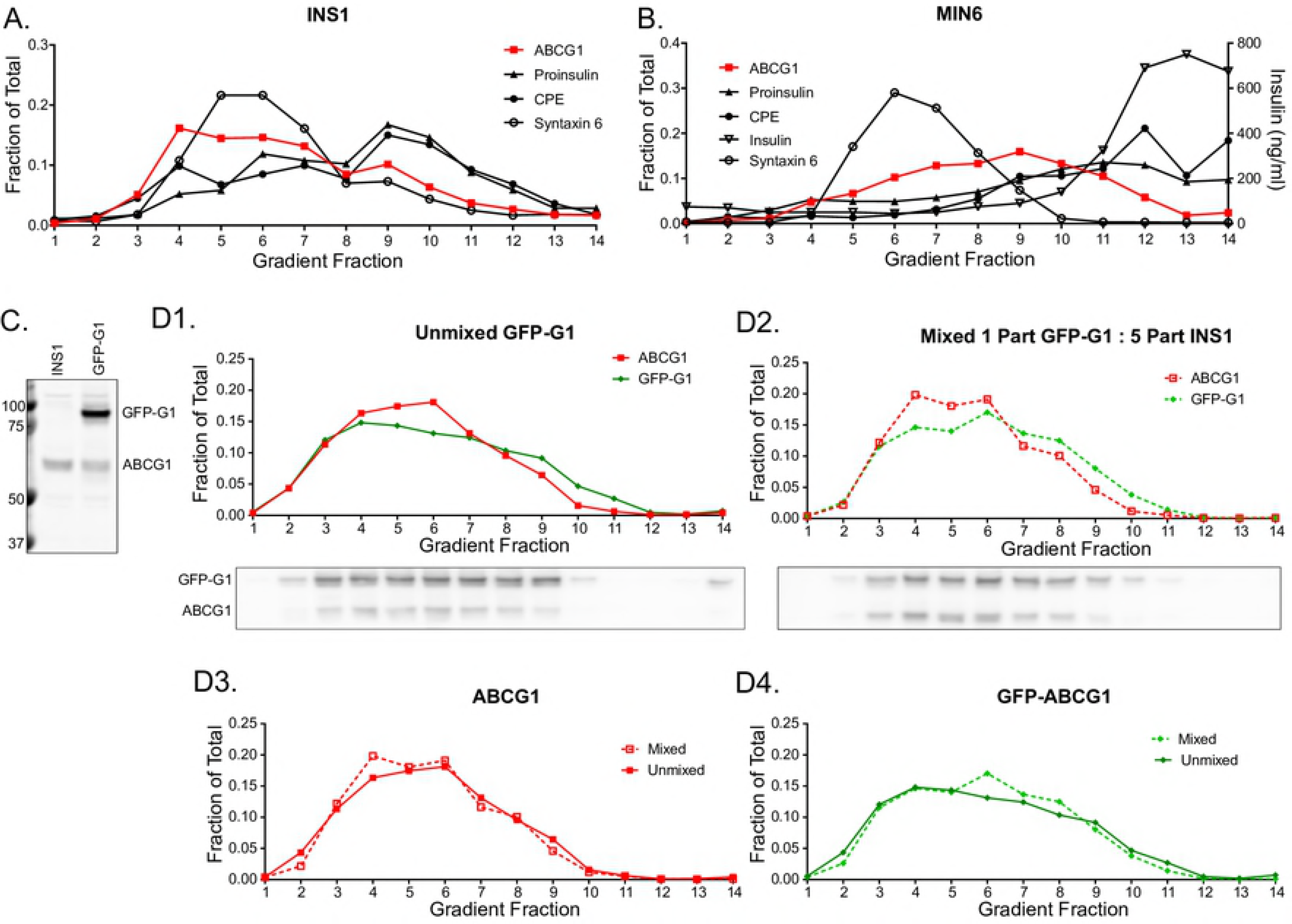
Distribution of ABCG1 by Cell Fractionation. (A) Fractionation of postnuclear supernatant from INS1 cells by centrifugation on a continuous sucrose density gradient showing the broad distribution of ABCG1 yet only modest overlap with secretory granule containing fractions. Marker distributions were quantified from western blots and are expressed as fraction of total detected on the gradients. Markers used are proinsulin (ER, TGN and immature granules), carboxypeptidase E (CPE; TGN, immature and mature granules), and syntaxin 6 (TGN and endosomes). (B) Fractionation of postnuclear supernatant from MIN6 cells confirming both the broad distribution of ABCG1, appreciable overlap with proinsulin-containing fractions (as reported previously [16]), and limited distribution in mature granules identified by insulin ELISA. Markers are as in (A). (C) Western blot using anti-ABCG1 showing the stable expression in INS1 cells of GFP-ABCG1 fusion protein (GFP-G1) at ~2.5-fold above the level of endogenous ABCG1. (D) Mixing experiment as analyzed by continuous sucrose gradient centrifugation of postnuclear supernatant fractions. Data are presented as fraction of total detected from western blots. Sample blots shown below the first two panels. Panel 1. GFP-G1 and endogenous ABCG1 distributions for cells stably expressing GFP-G1. Panel 2. Distributions observed for a mixed postnuclear supernatant containing 1 part GFP-G1 expressing cells and 5 parts non-transfected INS1 cells highlighting the very similar distributions of endogenous ABCG1 deriving mostly from the non-transfected cells and GFP-G1 from the transfected cells. Panel 3. Replots of endogenous ABCG1 distribution from panel 1 (Unmixed GFP-G1) and endogenous ABCG1 from panel 2 (Mixed PNS) illustrating that expression of GFP-G1 minimally alters the distribution of endogenous ABCG1. Panel 4. Replots of GFP-G1 distribution from panels 1 and 2 used as an index of reproducibility of the two gradients. Data presented are from one of two experiments with the same outcome.

Our new insight that ABCG1 did not co-fractionate with mature insulin granules also caused us to question our previously reported extensive co-localization of ABCG1 and insulin observed by immunofluorescence [16]. While we thought we had done appropriate controls to confirm specific labeling in the earlier study (which were presented in the Appendix), we felt compelled to reexamine this issue, and we discovered two problems, which together argued that it is necessary to revise our interpretation of the data. First, the fluorescently-labeled antirabbit secondary antibody that was used to detect rabbit anti-ABCG1 antibody was not sufficiently cross-adsorbed to fully abrogate its interaction with guinea pig anti-insulin antibody. Second, even though rabbit anti-ABCG1 antibody has been used for immunofluorescent localizations in macrophages and transfected cells where ABCG1 is relatively abundant [21,28], the low level of ABCG1 present in insulinoma cells necessitated using higher concentrations of both primary and secondary antibodies. As a consequence, fluorescent anti-rabbit antibody was binding sufficiently to guinea pig anti-insulin to report a false colocalization. In order to go beyond this problem, we decided to characterize the distribution of fluorescently-tagged ABCG1 (GFP-G1) that had been stably expressed in INS1 cells. Using western blotting, we determined that the cells were expressing the tagged transporter at ~2.5-fold above the endogenous level and that levels of free GFP or other lower molecular weight species were negligible (Fig 1C). We note that while use of CRISPR-Cas9 technology to tag endogenous ABCG1 would be a preferred approach for analyzing the distribution of ABCG1, we have been unsuccessful in developing a recombinant INS1 cell line for this purpose.

### GFP-G1 exhibits nearly the same distribution as endogenous ABCG1 and does not perturb the distribution of the endogenous transporter

To evaluate the extent to which GFP-G1 mimics the distribution of endogenous ABCG1 and also whether overexpression affects endogenous transporter distribution, we used cell fractionation and performed a mixing experiment. Parallel samples of GFP-G1 expressing cells and non-transfected INS1 cells were homogenized and spun at low speed to prepare postnuclear supernatants (PNS). On separate continuous sucrose gradients, we loaded PNS derived from GFP-G1 cells alone (unmixed) and a mixed PNS containing 1 part of GFP-G1 PNS and 5 parts of INS1 PNS (1:5 mix). In the 1:5 mix, ~84% of endogenous ABCG1 is contributed by the non-transfected INS1 cells and thus, to a very good approximation, reflects the distribution of the endogenous protein. After fractionation and western blotting using anti-ABCG1, the results (Fig 1D) indicate that: 1) the distributions of GFP-G1 and endogenous ABCG1 from the unmixed GFP-G1 PNS are very similar to each other, except that the proportions of GFP-G1 in fractions 5 and 6 and in fractions 9–11 are, respectively, slightly smaller and slightly larger than for endogenous ABCG1 (Fig 1D-1); 2) the distribution of GFP-G1 in the 1:5 mix is very similar to the distribution of endogenous ABCG1 in the 1:5 mix, although, the proportions of total GFP-G1 present in fractions 4–6 and in fractions 7–10 are, respectively, slightly smaller and slightly larger than the proportions of ABCG1 in the same fractions (Fig 1D-2); 3) the distributions of endogenous ABCG1 from the unmixed and 1:5 mix samples are very similar to each other (Fig 1D-3); and 4) GFP-G1 in the unmixed and 1:5 mix samples are practically identical (Fig 1D-4). Taken together, we conclude that GFP-G1 is reporting a distribution that is very similar to endogenous ABCG1, and that expression of GFP-G1 is not altering the distribution of endogenous ABCG1.

### GFP-G1 fluorescence co-localizes with markers of the TGN and ERC but not appreciably with mature insulin granules or markers of early and late endosomes

With the knowledge that exogenous GFP-G1 is a useful proxy for endogenous ABCG1, we used fluorescence microscopy to compare the localization of GFP-G1 to organelle markers detected by immunofluorescence in the transfected INS1 cells. GFP fluorescence was significantly accumulated perinuclearly where the signal appeared patterned but not tightly concentrated; occasional brighter puncta were visible at the pattern margins (Fig 2A). Comparison to immunostained insulin (Fig 2B) showed quite clearly that GFP-G1 did not mark intensely stained mature insulin granules that were present as multiple puncta mainly at the cell periphery. However, there was substantial overlap of GFP with the more diffuse perinuclear insulin staining, suggesting accumulation of ABCG1 at the TGN where insulin granule formation begins. Comparisons of GFP-G1 specifically to proinsulin (which is concentrated in the TGN and nascent granules) and to Golgin 97 (a *trans*-Golgi/TGN-associated tether protein) show quite similar signal patterns, supporting TGN localization (Fig 2C, D). Notably, the more intense foci of proinsulin staining were not fully matched by more intense foci of GFP fluorescence (inset in Fig 2C), suggesting that ABCG1 is not concentrated preferentially in forming granule membranes. As well, in the case of Golgin 97, GFP fluorescence shows a very similar pattern but is more broadly distributed and thus not tightly overlapping.

**Fig 2.**
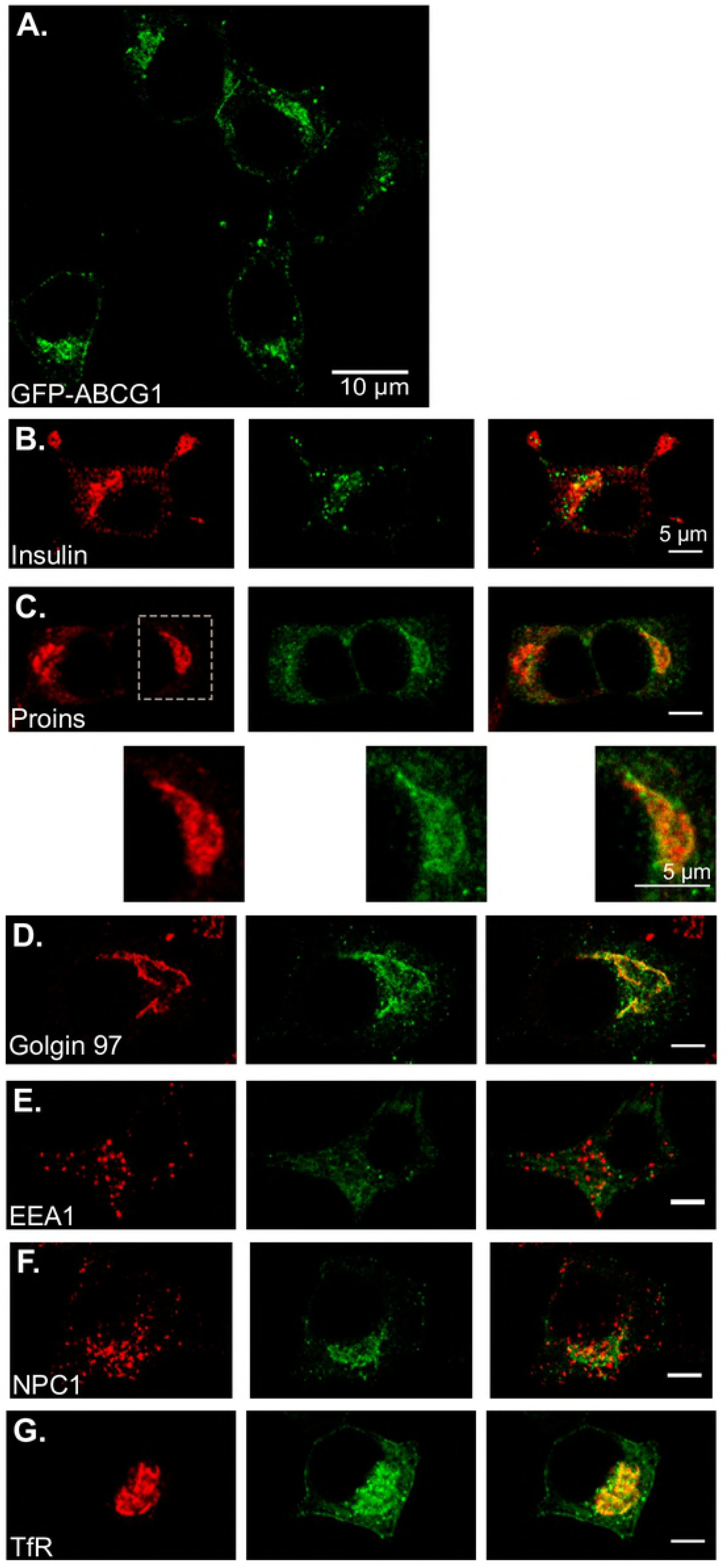
Distribution of GFP-G1 by confocal fluorescence microscopy as compared to organelle markers. (A) Low magnification survey illustrating that GFP-G1 generally is accumulated perinuclearly, mostly in a finely granular pattern with occasional surrounding brighter puncta. (B-G) Higher magnification images comparing the distribution of GFP-G1 to organelle markers. (B) Little co-localization with insulin at the cell periphery where mature granules accumulate but noticeable overlap perinuclearly where storage granules are being formed. (C) Extensive overlap with proinsulin accumulated perinuclearly and resembling the perinuclear overlap seen in panel B. Expanded insets (below) show the GFP-G1 fluorescence has the same pattern as proinsulin but is not especially co-concentrated with proinsulin in brighter puncta of the latter. (D) GFP-G1 exhibits a highly similar pattern as Golgin97 but is more diffusely distributed. (E) Little overlap of GFP-G1 with EEA1 marking early endosomes. (F) Quite modest overlap of GFP-G1 with NPC1 marking late endosomes and lysosomes. (G) Substantial overlap of GFP-G1 and exhibiting a similar pattern of distribution as transferrin receptor TfR concentrated in the endosomal recycling compartment. GFP-G1 is also visible at the plasma membrane and diffusely within the cytosol (presumably ER).

Because various endocytic compartments substantially accumulate in the perinuclear region and have been identified as sites where ABCG1 resides in other cell types [21,24,28], we compared the distribution of GFP-G1 to EEA1, an early endosomal marker; NPC1, a marker for late endosomes and lysosomes; and transferrin receptor (TfR), a marker for the endosomal recycling compartment (ERC). GFP fluorescence was minimally colocalized with EEA1 (Fig 2E) and NPC1 (Fig 2F) and also did not colocalize with the lysosomal membrane protein LIMP2 (unpublished data). However, in the case of TfR staining, GFP-G1 signal overlapped quite well (Fig 2G), similar to what we observed with proinsulin and Golgin 97. While the diffuse GFP fluorescence clearly extended beyond the more concentrated TfR staining, there were some focal regions of co-concentration (intense yellow in the merged image). Also, low levels of GFP-G1 can be seen at the plasma membrane as well as diffusely distributed in the cytoplasm, the latter consistent with detectable presence in the ER (e.g., Fig 2G). Plasma membrane staining is consistent with our previous measurements using cell surface biotinylation showing that ~13% of total endogenous ABCG1 is exposed on the surface of pancreatic MIN6 cells [16]. Detectable presence in the ER likely reflects continual synthesis of ABCG1, which undergoes turnover quite rapidly (see below).

In an effort to identify the occasional larger, bright GFP-G1-positive puncta that were mostly distributed peripheral to the perinuclear staining, we examined possible association with autophagosomes using a fluorescently-tagged marker LC3-mRFP. Here, we observed colocalization and co-migration with a portion of the puncta (Fig 3A and Supplementary video 1). Also, we imaged live cells that had been incubated with the cathepsin D Magic Red substrate (a marker for cathepsin-active lysosome-related compartments) and saw colocalization and comigration with fewer of the bright puncta (unpublished data). Together, these observations suggest that at least some of the bright GFP-positive puncta may be autophagosomes and autolysosomes. Because these puncta are variably present but generally more noticeable in cells expressing higher levels of GFP-G1, we surmise that their presence may be related to overexpression. Thus, they may not be on the main trafficking pathways of ABCG1. Interestingly, live imaging additionally showed that there was a very robust trafficking toward and away from the cell periphery of bright GFP-G1-positive puncta that were mainly smaller than those labeled by LC3 emphasizing the itinerant behavior of the transporter (Fig 3B and Supplementary video 2).

**Fig 3.**
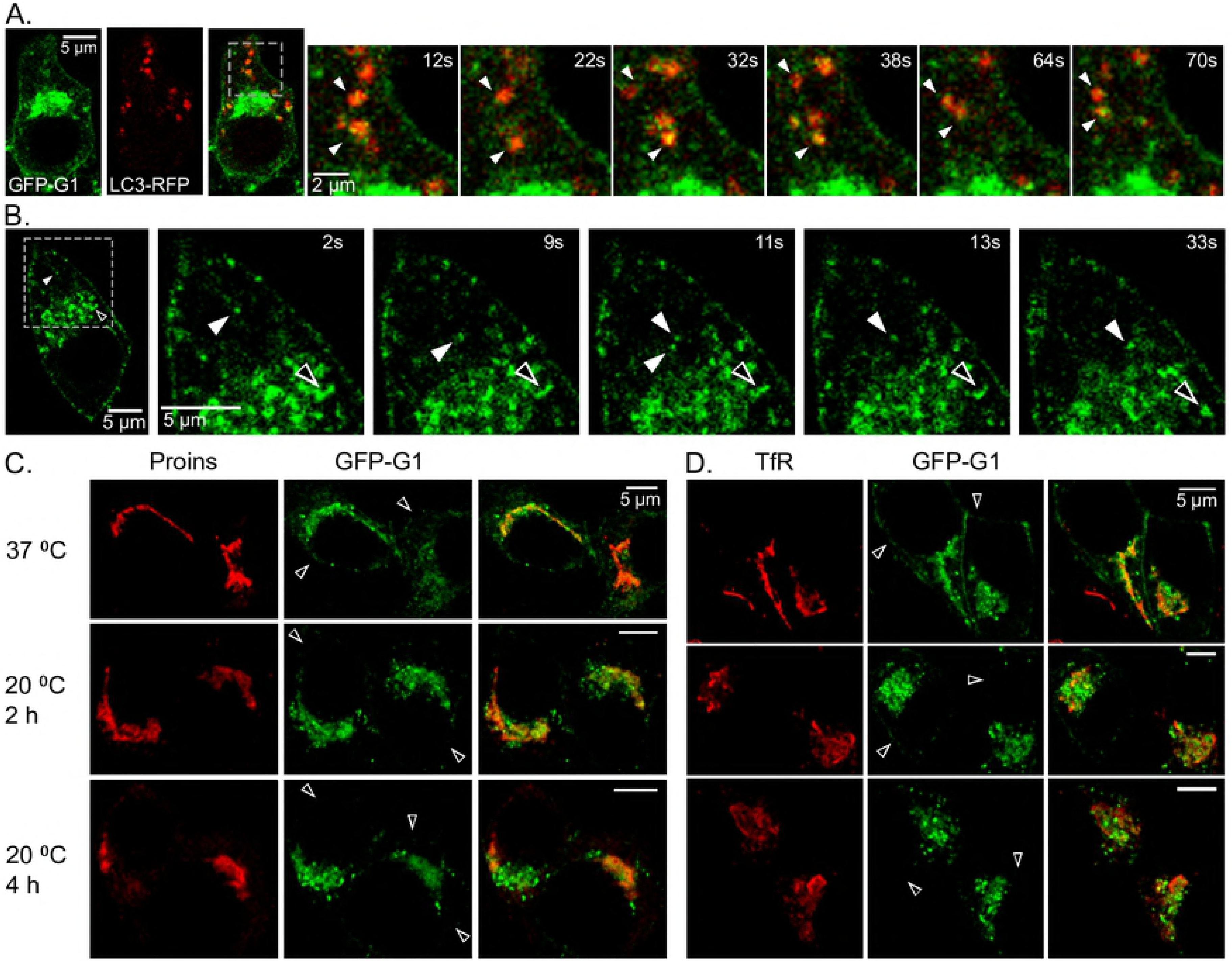
Manipulating and visualizing GFP-G1’s dynamic distribution. (A) Association of autophagosomal marker LC3-mRFP with a subset of the brightly fluorescent puncta of GFP-G1 shown by live imaging. Lower magnification shows overlap in the peripheral cytoplasm and higher magnification is a time series captured from the indicated region. Frames are taken from Supplementary Video 1 and have been selected to illustrate colocalization and co-migration of LC3 and GFP-G1; arrowheads highlight specific examples. (B) Series of frames from live imaging (from Supplementary Video 2) selected to highlight active trafficking of small GFP-G1 puncta (arrowheads). Video illustrates bi-directional movement toward and away from the cell periphery. (C) Incubation at 20 °C increases perinuclear accumulation of bright GFP-G1-positive puncta adjacent to where proinsulin immunostaining overlaps diffuse GFP-G1 fluorescence. (D) Increased perinuclear accumulation of GFP-G1 at 20 °C also clusters near and partially overlaps TfR immunostaining. In both C, D, peripheral cell surface GFP-G1 fluorescence is greatly decreased by the low-temperature incubation (arrowheads).

### Further insights into the colocalization of GFP-G1 with the TGN and ERC

We carried out additional studies to evaluate the accumulation of GFP-G1 in the TGN and ERC. First, we subjected cells to incubation at 20 °C prior to fixation and immunostaining for proinsulin and for TfR. This strategy is well known to impose a block to membrane trafficking from the TGN [29,30] and the ERC [31]. At 2h and 4h incubation at 20 °C, GFP-G1 progressively became more highly concentrated in perinuclear puncta that were closely apposed to both proinsulin and TfR. Where GFP-G1 directly overlapped the markers, it remained more diffuse (Fig 3C, D). Over the same timeframe, the signal at the plasma membrane was substantially depleted (Fig 3C, D). Together, these features are consistent with the presence of GFP-G1 in both the TGN and ERC but also strong accumulation in trafficking carriers that are hindered in their fusion and/or trafficking along secretory and recycling pathways due to the decreased temperature. Also, we noted, particularly at 4h, that diffuse cytoplasmic staining for proinsulin became more apparent (Fig 3C), which is consistent with slowed exit and increased accumulation in the ER at the lower temperature.

### ABCG1 has a short half-life in INS1 cells and is degraded by both ERAD and lysosomes

The accumulation of GFP-G1 staining in multiple cellular compartments, its robust trafficking, and its generally diffuse appearance at these sites suggested that ABCG1 might not have a specific compartmental residence. Rather, since ABCG1 is known to be a short-lived protein in other cell types [32–34], we wondered whether its distribution might reflect its brief lifetime. Thus, upon exit from the ER and transit through the Golgi, it would pass onward through the cell’s constitutive trafficking pathways exhibiting accumulation at sites (e.g., TGN and ERC) where membrane exit is rate-limiting. To determine whether ABCG1 is indeed shortlived in INS1 cells, we used pulse labeling with ^35^S-amino acids and tracked the progressive loss of radiolabeled ABCG1 during chase incubation by immunoadsorption with anti-ABCGI and phosphorimaging. As shown in Fig 4A, turnover is indeed quite rapid with a t_1/2_ of < 2h. To determine where ABCG1 was being turned over, we carried out a follow-up study in which we treated cells with cycloheximide to block protein synthesis and examined how the ensuing transporter loss was affected by MG132 (a proteasomal inhibitor that diagnoses membrane protein loss by ER-associated degradation ‑ ERAD), bafilomycin A (a proton pump inhibitor that diagnoses lysosomal degradation), or a combination of the two inhibitors. MG132 and bafilomycin A each slowed the rate of loss with MG132 having the larger effect, and the combined inhibitors almost completely blocked loss (Fig 4B). Thus, a substantial portion of ABCG1 in INS1 cells is removed from the ER by ERAD while the remainder passes constitutively along transport routes to the TGN, cell surface, endosomes, and ultimately to degradation in lysosomes.

**Fig 4.**
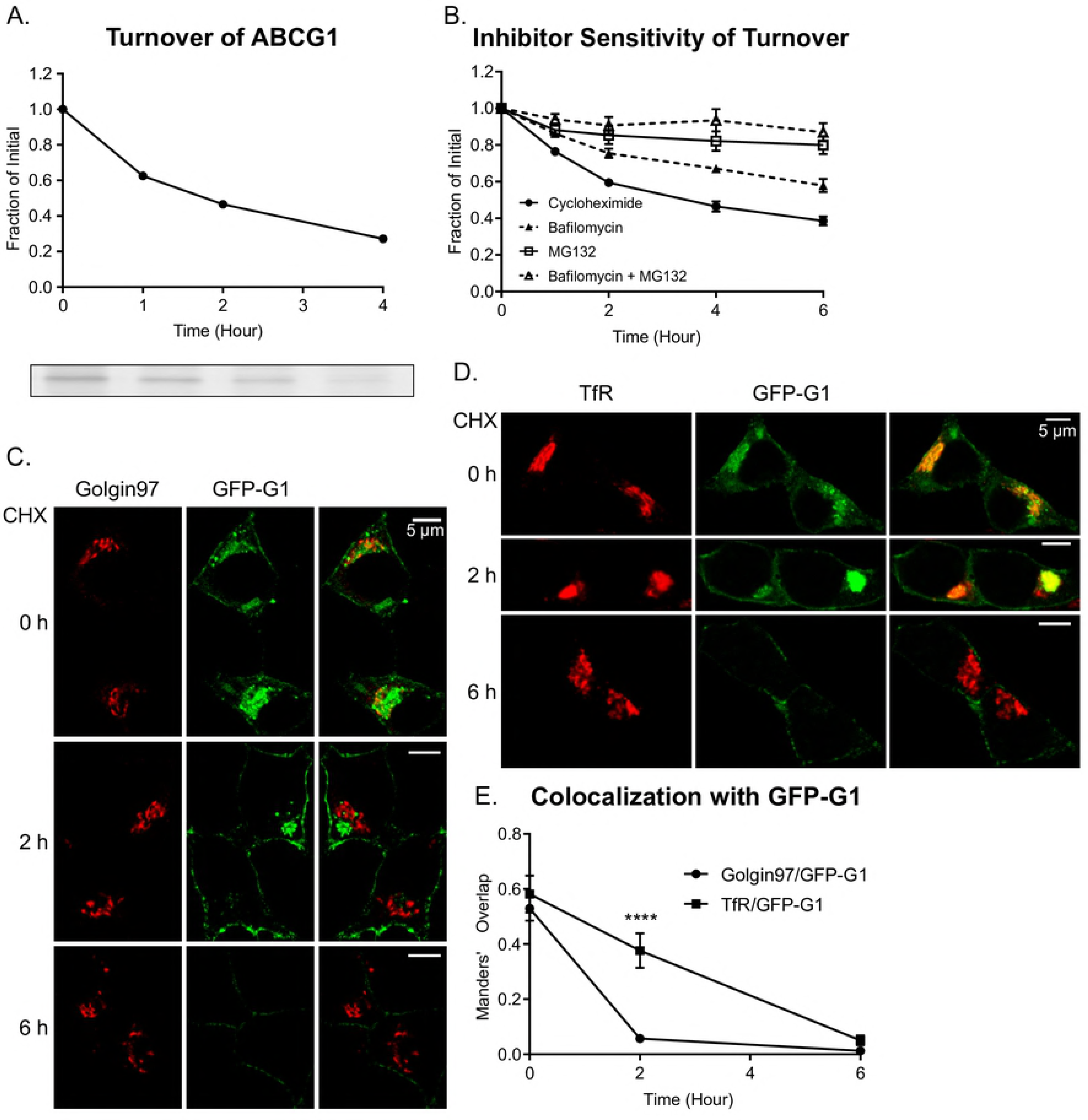
Turnover of ABCG1 and GFP-G1. (A) Pulse-chase biosynthetic labeling with ^35^S-amino acids and immunoadsorption of ABCG1. Phosphorimage (below) and quantification (above) show that the transporter is short-lived in INS1 cells with a halftime of turnover of < 2h. (B) Results from western blotting showing loss of ABCG1 following inhibition of protein synthesis (by cycloheximide) occurs by both ERAD/proteasomal degradation (inhibited by MG132) and lysosomal degradation (inhibited by the acidification blocker bafilomycin A). Data are from 7 experiments; error bars indicate SEM. (C) Representative fluorescence images from samples treated for 0, 2, and 6h with cycloheximide (CHX). Both diffuse cytoplasmic GFP-G1 (presumed to be ER associated) and GFP-G1 colocalized with the TGN marker Golgin 97 are lost by 2h while the remainder of accumulated perinuclear GFP-G1 and much of the cell surface transporter is lost by 6h. (D) Complementary images to panel C showing perinuclear GFP-G1 remains colocalized with TfR at 2h post cycloheximide addition but is lost by 6h. (E) Colocalization presented as Manders’ coefficients comparing overlaps of Golgin 97 and TfR with GFP-G1 during cycloheximide treatment. Error bars are SEM. ****, p < 0.0001. Detailed cell-by-cell scatter plots shown in Figure S2.

To complement the kinetic measurements of ABCG1’s lifetime, we followed the changes in the amount and localization of GFP-G1 by confocal fluorescence microscopy after treating cells with cycloheximide. To provide intracellular landmarks, we immunostained the specimens with Golgin97 (TGN marker) and TfR (ERC marker). As can be seen in Fig 4C, there are readily detectable changes in the initial distribution of GFP-G1 by 2h and 6h of cycloheximide treatment. Both the diffuse cytoplasmic GFP fluorescence, presumed to be ER, and the fluorescence that is colocalized with Golgin97 disappear by 2h while fluorescence that is colocalized with TfR as well as signal at the plasma membrane remain readily visible (Fig 4D). Rapid loss from the ER is consistent with depletion by both ERAD and transport to the Golgi, while depletion from the TGN reflects combined loss of input from ER/Golgi and trafficking along post-TGN pathways. By 6h, the fluorescence colocalizing with TfR largely disappears and cell surface staining is decreased, consistent with cycling to and from the plasma membrane and progressive trafficking to lysosomes. The residual staining at the cell surface is similar to observations made for heterologous expression of ABCG1-GFP in HeLa cells [24]. These results, which are quantified in Fig 4E and Fig S2, clearly highlight the itinerant behavior and expeditious turnover of ABCG1 and clarify its transient accumulation in both the TGN and the ERC during its transit through the cell.

### Effects of ABCG1 depletion on endocytic trafficking and transferrin receptor levels

Earlier studies have shown that the ERC normally is a significant site of intracellular cholesterol accumulation [35]. Because we have observed that GFP-G1 accumulates in the ERC, we sought to detect whether ABCG1 might contribute to the functional organization of the ERC. Thus, we used siRNA-mediated knockdown to reduce endogenous ABCG1 by ~90% in INS1 cells, an approach whose specificity and on-target actions we recently validated [27], and then followed the trafficking of fluorescent transferrin and of fluorescent cholera toxin B subunit (CTxB) in comparison to control knockdown cells. Transferrin (Tf) is well known to undergo receptor-mediated, clathrin-dependent endocytosis and to accumulate in part in the ERC along its recycling pathways back to the plasma membrane [26], whereas CTxB (like shiga toxin B subunit) bound to the cell surface is internalized substantially by clathrin-independent endocytosis and co-accumulates with Tf in the ERC before retrograde transport from the ERC to the Golgi [36–38]. Notably, the transit of CTxB to the Golgi via the ERC is also followed by chimeric carboxypeptidase E-Tac, and may trace a recycling route for selected neuroendocrine granule membrane proteins [39].

We carried out separate sets of experiments to examine the recycling of endocytosed fluorescent Tf from the ERC to the plasma membrane and the internalization and subsequent trafficking of both Tf and CTxB. To analyze Tf recycling, we preloaded the cells that had been plated on coverslips with rat Tf-Dylight488 at 37 °C, switched to fresh medium containing unlabeled Tf and at specified timepoints, washed, fixed and surface-labeled the cells as indicated under Methods. From the captured fluorescence microscopic images, total Tf fluorescence per cell area (defined by surface labeling with fluorescent wheat germ agglutinin) was quantified for multiple cells in multiple fields to enable tracking of progressive loss of fluorescence as Tf recycled (See Methods). Sample images taken at 5 and 40 min in control and ABCG1-depleted cells are shown and the efflux of fluorescence is plotted in Fig 5A. As can be seen, the overall cellular content of fluorescent Tf is significantly higher initially in ABCG1 knockdown cells. The ensuing efflux, however, occurs at a comparable rate (fractional decrease of mean intensity) in the two types of samples. Thus, ABCG1 deficiency increases the accumulation of Tf in the ERC, but it does not appear to perturb recycling.

**Fig 5.**
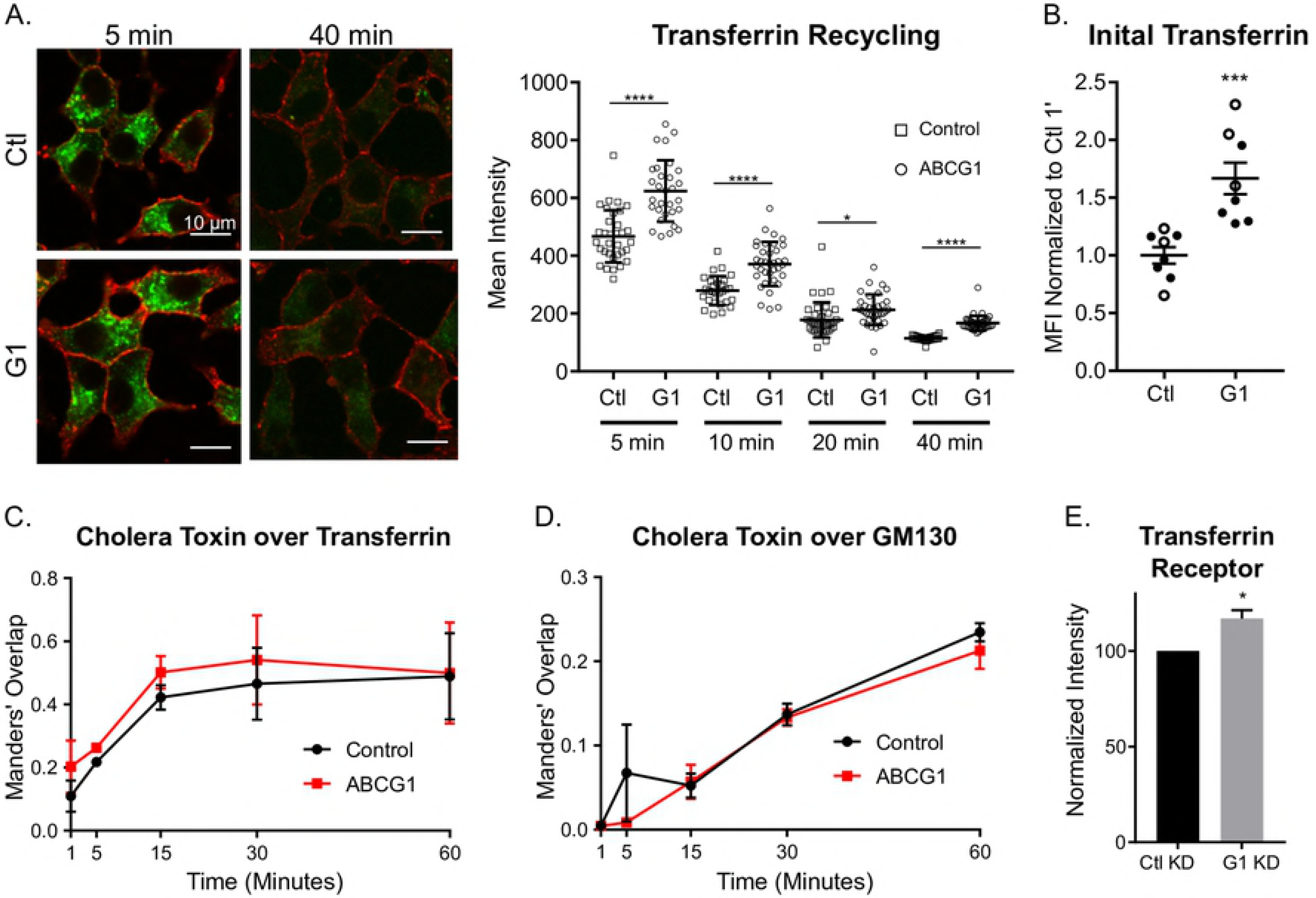
ABCG1 depletion does not noticeably perturb membrane trafficking through the ERC but significantly increases the level of transferrin receptor. (A) Representative images taken at 5 and 40 min following preloading of cells with Dylight-488 rat transferrin (Tf) illustrating recycling of Tf from control and ABCG1-depleted cells. Cell perimeters were labeled with fluorescent wheat germ agglutinin. Loss of ABCG1 does not significantly perturb recycling, although transporter-depleted cells accumulate more fluorescent Tf during loading. The accompanying graph compares efflux during the recycling incubation and is plotted as mean intensity of Tf fluorescence per cell profile area. At the outset and throughout recycling, Tf in ABCG1-depleted cells is significantly higher than in Control (* p<0.05; **** p<0.0001). However, fractional disappearance of Tf between successive timepoints is about the same for both samples (~38%). (B) Fluorescent Tf bound and newly internalized is notably increased in ABCG1-depleted cells; ***, p<0.001. Data are normalized to the mean of 1-min control in each of two experiments. (C) Colocalization of CTxB and Tf increases and plateaus at the same rate in control and ABCG1-depleted cells. (D) Progressive increase in colocalization of CTxB and GM130 shows that retrograde trafficking to the Golgi does not differ significantly in control and ABCG1-deficient cells. Data in (C) and (D) are presented as Manders’ coefficients, determined cell-by-cell at each timepoint. (E) Level of TfR in control and ABCG1 KD cells determined by western blotting in four independent experiments; *, p<0.05.

To compare uptake and subsequent trafficking of probes in control and ABCG1-knockdown samples, cells plated on coverslips were chilled to 0 °C and surface labeled using 10 μg/ml rat Tf-Dylight488 and 20 nM CTxB-Alexa555, washed and then warmed to 37 °C for increasing times up to 60 min. At each timepoint, the samples were rinsed in PBS, fixed, immunostained for the Golgi marker GM130 and examined by confocal fluorescence microscopy (see Methods). In general, we found that the uptake and concentration of the probes in the ERC was slower than in BSC-1 or HeLa cells (where concentration in the perinuclear ERC was extensive by 7 min [36]). We also observed that the level of cell-associated CTxB-Alexa555 (but not of Tf-Dylight488) varied over a wide range (Fig S4A). For quantitative comparison of control and ABCG1-depeleted cells, we measured: 1) Tf intensity per cell area initially (1 min) to estimate levels at (and newly internalized from) the cell surface; 2) the progressive colocalization of Tf and CTxB during their convergence at the ERC; and 3) the progressive colocalization of CTxB and GM130 to assess retrograde trafficking to the Golgi. Interestingly, we observed that initial bound and internalized Tf was significantly higher in ABCG1-depleted cells as compared to the control, suggesting that surface levels of TfR are elevated (Fig 5B). Regarding convergence of Tf and CTxB at the ERC, the colocalization of the two endocytic probes appeared to progress more quickly in ABCG1-depleted cells (Fig 5C). However, the difference did not reach significance, and the increased overlap in the knockdown may reflect, at least in part, the higher initial Tf level (Fig 5B). In the case of retrograde trafficking of CTxB to the Golgi, overlap of CTxB and GM130 progressively increased beyond 5 min, and the rate did not differ significantly between control and ABCG1 KD samples (Fig 5D). Sample images comparing the localizations of Tf and CTxB and of CTxB and GM130 at 5, 15, and 30 min are shown in Fig S4A.

Our observations that fluorescent Tf accumulation in the ERC at the outset of recycling analysis and early appearance at the cell periphery during uptake analysis were both elevated in ABCG1-deficient cells suggest altered TfR trafficking but also could result from an increased receptor level in the knockdown cells. To investigate this, we examined the total amount of TfR in samples of cell lysate by western blotting. As shown in Fig 5E, loss of ABCG1 indeed increased TfR by ~20%. Finally, to check whether there might be noticeable changes in the organization of endosomal recycling pathways as a result of ABCG1 depletion, we compared the localizations of EEA1 and TfR by immunofluorescence in control and ABCG1 KD cells. For this study, we visually compared the sizes of EEA1-stained puncta and also searched for any clear differences in the distribution of TfR in the cytoplasm using Golgi α-mannosidase-II immunostaining as a well-defined perinuclear point of reference. In neither case did we observe changes in the ABCG1-deficient cells (sample images shown in Fig S4B).

These initial efforts to explore endocytic processes in ABCG1-depleted cells have not identified major consequences of transporter deficiency to the standard operation of constitutive recycling and retrograde trafficking. However, knockdown significantly increases cellular TfR levels, which may reflect either progressively increased expression or decreased degradation of the receptor as transporter levels decrease. While at this point, we have not searched more broadly for other possible perturbations that involve endocytic trafficking and the ERC in particular, we recognize from our earlier studies of proinsulin transport and insulin storage that the effects of ABCG1 loss are both partial and manifest in selected post-Golgi pathways [27]. The same may apply to endosomal sites that accumulate ABCG1. Regardless, our present results point to likely roles of ABCG1 beyond the site of insulin granule formation in INS1 cells.

## Discussion

In this study, our main mission has been to reevaluate and clarify the localization, trafficking, and sites of function of ABCG1 in pancreatic beta cells. At the outset, we realized from cell fractionation studies using insulin-secreting cell lines that ABCG1 did not co-localize appreciably with markers of insulin granules. Thus, we were compelled to modify our previous view [16]. Indeed, our current results indicate that little ABCG1 is associated with insulin granules, an observation that seems consistent with the short half-life of the transporter that we now report. We note, however, that we have not fully ruled out its presence at low levels in granule membranes, especially in islet beta cells that are more differentiated and maintain a much larger granule storage pool. Indeed, as mentioned under Results, our earlier purification of insulin granules from pancreatic islets indicated presence of ABCG1 in lower density granule fractions [16]. Consequently, it remains possible that ABCG1 maintains association with a subset of granules, particularly during their maturation. This possibility is consistent with our recent demonstration that ABCG1 supports the process of insulin granule formation [27]. Nevertheless, based on our present results, the majority of ABCG1 is clearly not granule-associated.

Despite extensive effort, we have been unsuccessful in identifying antibodies suitable for immunolocalization of ABCG1 in our cultured cells. We therefore examined the distribution of GFP-tagged ABCG1 stably expressed in INS1 cells instead. We validated this approach using mixing experiments based on cell fractionation and showed that GFP-G1 is an appropriate proxy for endogenous ABCG1. In the ensuing analysis of GFP-G1 distribution by fluorescence microscopy, we noted that most of the transporter is not highly concentrated in the manner typically observed for organelle markers. While GFP-G1 accumulates in patterns that are closely similar to those of TGN-and ERC-associated marker proteins (Fig 2), its fluorescence remains diffuse. This appearance does not seem to reflect changes incurred during fixation and permeabilization preceding examination, as the same diffuse accumulation is observed when imaging live cells (Fig 3A, B). Notably, our observation that GFP-G1 fluorescence does not appreciably co-concentrate with proinsulin, even though it is in close proximity, suggests that as ABCG1 aids in generating the cholesterol-enriched granule membrane [27], it may be progressively excluded and thus traffics along other TGN-and immature granule-derived pathways leading to the cell surface and to endosomes [40,41]. These same pathways either directly or indirectly lead to the ERC and thus to ABCG1’s accumulation there. But in the ERC, its appearance remains diffuse, arguing that it is not a persistent ERC resident. Thus, ABCG1 acts as an itinerant protein that accumulates where there are slow steps in intracellular transport. The tendency of ABCG1 to not associate with cholesterol-enriched domains such as those thought to be generated in forming insulin granules and present in the ERC seems consistent with the previous observation that the transporter facilitates transbilayer translocation of cholesterol within less-ordered domains [8].

Our efforts to localize GFP-G1 also indicate its presence on the plasma membrane of INS1 cells, which is in agreement with our previous assessment in MIN6 cells [16]. The slow clearance of surface-associated GFP-G1 (Fig 3C) and the slower clearance of cell surface (as compared to diffuse intracellular) fluorescence during cycloheximide treatment (Fig 4) suggest that ABCG1 in β cells may form some sort of stabilized interaction at the cell surface. The punctate staining along the plasma membrane (Fig 2G, 3B) and persistence there late after protein synthesis inhibition (Fig 4C, D) may reflect associations of ABCG1 either with the actin cytoskeleton as recently reported for fluorescently-tagged ABCG1 expressed in CHO and HeLa cells [42] or membrane lipid microdomains (e.g., as at least a portion of ABCG1 is known to be palmitoylated [43]). Moreover, sustained residence at the cell surface may also be indicative ofa lipid-regulatory function at this site. Nevertheless, GFP-labeled plasma membrane-derived endocytic vesicles that we readily detected by live cell imaging (Fig 3B and Supplementary video 2) argue that there is robust trafficking of ABCG1 into and out of the cell surface as would be anticipated for an itinerant protein.

In analyzing the turnover of ABCG1 in INS1 cells, our studies using MG132 and bafilomycin A imply that degradation occurs primarily through ERAD/proteasomes (as reported elsewhere [34]) while lysosomal degradation accounts for the remaining loss. We found that turnover was not affected by the calpain inhibitor calpeptin (unpublished data); while calpain has been implicated in degrading a portion of the functionally related ABC transporter ABCA1 [44,45], it does not appear to be a factor in ABCG1 turnover in insulin secreting cells or elsewhere [34]. Interestingly, our observation that lysosomes play a significant role in the turnover of ABCG1 differs from observations in both macrophages (for endogenous ABCG1) and CHO cells (engineered to stably express ABCG1) where the bulk of turnover appeared to be mediated by proteasomes and thus presumably ERAD [34]. We surmise that the difference could reflect the distinct cell type and hence range of potential roles of ABCG1 in β cells. Loss of more than half of ABCG1 by ERAD/proteasomal degradation is consistent with the rapid loss of diffuse cytoplasmic staining of GFP-G1 during cycloheximide treatment (Fig 4F) and is similar to what has been observed for the cystic fibrosis transmembrane conductance regulator CFTR, another ABC transporter, using other approaches [46]. Taking this rapid loss into account, the further disappearance of much of the rest of endogenous ABCG1 and GFP-G1 (Fig 4) by 4–6h indicates that the remaining transporter is also not long-lived and is being lost at a rate that is quite rapid compared to recycling cell surface receptors such as TfR and the LDL receptor (Fig 4 versus [47,48]). Thus, while ABCG1 is transported at appreciable levels in the β cell’s secretory and endocytic pathways, we suspect that its trafficking may be biased by sorting or modifications (e.g., ubiquitylation [49]) that favor lysosomal degradation over constitutive recycling.

Another realization from these studies is that ABCG1 probably functions at multiple locations within the cell, possibly even beyond the sites where it transiently accumulates. This would not be surprising if ABCG1’s mechanism of action involves transmembrane translocation of phospholipids [50,51] and consequent redistribution of cholesterol and other non-polar lipids to reestablish their equal chemical potentials within bilayer leaflets as hypothesized [8]. While testing this role experimentally is an important goal, we note that current approaches to evaluating cholesterol redistribution, e.g., by altered filipin staining, do not have sufficient sensitivity and resolution to detect what are likely to be relatively small changes in concentration, in particular perinuclear compartments [27]. In principle, however, ABCG1 may carry out its role in any compartment beyond the ER, i.e., where it is dimeric and thus active. If this is the case, then the various reports that ABCG1 is acting in different compartments such as late endosomes [24,28], the plasma membrane [2,8,23,52], and the TGN [27] are not contradictory. Differences in localization/accumulation and reported function may largely relate to the cell type being examined and the membrane trafficking kinetics in that cell. Moreover, our findings here and elsewhere [27] make the case that ABCG1’s action in INS1 cells is manifest in the TGN and elsewhere, possibly including other well-recognized sites of cholesterol accumulation and potential redistribution [35]. We propose that ABCG1 might best be viewed as a roving sensor and modulator of membrane biophysical properties that in beta cells is tuned to the function of the insulin secretory pathway. Its constitutive synthesis, robust trafficking and rapid turnover by both ERAD and lysosomal degradation provide mechanisms for maintaining transporter levels that are optimized for the level of proinsulin biosynthesis/transport and the related requirements for packaging and storing insulin in stable cholesterol-enriched membranes. In the broader context, these same characteristics of ABCG1 seem especially well suited for a transporter that is potentially active in all post-Golgi compartments and where local accumulation, such as in reverse cholesterol transport, alters cholesterol distribution and/or the balance of membrane order vs. disorder. Indeed, ABCG1 may be performing functions that are complementary to those deduced for ABCA1 not only in reverse cholesterol transport but also in ABCA1’s role in adjusting biophysical properties of the plasma membrane to accommodate to the local environment within cell communities or to regulate inflammatory responses [10,11]. Finally, whether ABCG1 is functioning at the plasma membrane or at multiple sites in its more global regulation of cell proliferation or differentiation [12–14,53] remains to be determined.

## Materials & Methods

### Reagents

Rat fluorescent transferrin (Tf-DyLight488) was from Jackson Immunoresearch. Fluorescent cholera toxin B-subunit (CTxB-Alexa555 or 568) was from Invitrogen/Molecular Probes. Wheat germ agglutinin-Alexa555 was from Thermo-Fisher. Unlabeled rat transferrin was from Rockland Scientific. Cycloheximide, and bafilomycin A were obtained from Sigma/Aldrich, MG132 was from Cayman Chemical Co., calpeptin from Calbiochem, Magic-Red cathepsin D substrate from Neuromics.

### Cells and Culture

INS1 823/13 cells [54] were maintained in RPMI 1640 medium, 10 mM Hepes, 1 mM sodium pyruvate, 50 μM β-mercaptoethanol, 1x pen/strep, and 10% FBS (Atlanta Biological). MIN6 cells were a gift from Dr. Chien Li (University of Virginia and NovoNordisk) and were maintained in DMEM containing 15% FBS (Atlanta Biological) and 1x pen/strep.

### Plasmids and expression

Mouse ABCG1 cDNA in pcDNA3.1 (obtained from the Hedrick Laboratory, LaJolla Institute of Allergy and Immunology) was used to generate N-terminally EGFP-tagged ABCG1 (GFP-G1) by insertion into pmEGFP_C1 (Invitrogen/ThermoFisher). The sequence was confirmed and the construct was transfected into INS1 823/13 cells using Amaxa. Expressing cells were selected for neomycin resistance and small cell clusters (typical of endocrine cell growth) were isolated for subsequent propagation using cloning rings. Selection and enrichment of expressing cells were validated by fluorescence microscopy, and cells were maintained as for parent INS1 cells except in the continued presence of neomycin. A construct encoding LC3-mRFP in pcDNA3.1 was the kind gift of James Casanova, and was transiently expressed in GFP-G1 cells using Lipofectamine 2000.

### Antibodies used for western blotting, immunofluorescence and immunoprecipitation

Antibodies to the following proteins were used in this study: Porcine insulin, polyclonal guinea pig, Dako A0564; rat proinsulin for immunofluorescence, mouse Mab, Alpco (gift, 5 mg/ml; used at 1 μg/ml); proinsulin for western blotting, rabbit polyclonal, Santa Cruz sc-9168; CPE, mouse Mab, BD Biosciences 610758; EEA1, mouse Mab, BD Biosciences 610457; NPC1, rabbit Mab, Epitomics EPR5209; Transferrin Receptor, mouse Mab 68.4 ATCC; ABCG1 used for immunoprecipitation (2 μl per sample), rabbit polyclonal, Novus Biologicals #NB400-132 Lot E; Syntaxin 6, mouse Mab 3D10, Stressgen ADI-VAM-SV025D; Actin, polyclonal rabbit Cytoskeleton, AAN01, lot 45; α-mannosidase-II, polyclonal rabbit, gift of Kelley Moremen, University of Georgia.

For anti-ABCG1, commercial antibodies did not perform well on pancreatic cells containing relatively low amounts of antigen compared to macrophages or cells overexpressing exogenous ABCG1. Therefore, as recently described [27], we developed an antibody to the synthetic peptide (C)KKVDNNFTEAQRFSSLPRR-NH_2_ (within the N-terminal cytoplasmic domain of ABCG1). The antibody was made by Pacific Immunology Corp and was affinity purified using peptide coupled to Sulfolink (Pierce/ThermoFisher) via the appended N-terminal cysteine. The antibody was deemed not satisfactory for immunofluorescence but showed excellent specificity in western blots comparing postnuclear supernatant fractions from control and ABCG1-deficient cells (Fig S1 in [27]). Antibody was used at a final concentration of 0.2 μg/ml.

Secondary antibodies for immunofluorescence were Alexa-conjugated from Molecular Probes/Life Technologies and Jackson Immunoresearch (the latter for guinea pig cross-adsorbed). When pairing other antibodies with guinea pig anti-insulin, use of secondary antibodies cross-adsorbed against guinea pig was essential. Secondary antibodies for western blotting were from Licor: Goat anti-Rabbit 696 and Goat anti-Mouse 800.

### Continuous Sucrose Gradients

The distributions of endogenous ABCG1, stably expressed GFP-ABCG1 and a variety of marker proteins were compared by cell fractionation using centrifugation of postnuclear supernatants on continuous sucrose gradients. Cells (INS1 or MIN6) scraped and suspended in 0. 29 M sucrose, 5 mM MOPS, 0.2 mM EDTA (pH 7.2) supplemented with protease inhibitor cocktail (Complete Mini, Roche) were homogenized using a ball bearing homogenizer (0.2507in cylinder; [55]) 8 passes with 0.2496-in ball bearing. The homogenate was spun 1000 x g, 10 min in a microcentrifuge to prepare a postnuclear (PNS) fraction. Aliquots of the PNS (including those prepared by mixing PNS from GFP-G1 expressing and control cells as described in the Results) were adjusted to 1 mM EDTA and loaded on continuous (0.6–1.6M) sucrose gradients prepared in Beckman SW41 tubes and spun overnight to equilibrium (≥ 18 hr, 32,000 rpm). Fractions collected manually were checked for refractive index (to confirm near identity of gradients being compared), diluted with 0.2 M MOPS, pelleted (40 min, 60,000 rpm) in a TLA120.2 rotor, resuspended in a measured volume of residual supernatant, and solubilized in sample buffer plus DTT. An equal portion of each gradient fraction was used for gel electrophoresis and western blotting enabling calculation and plotting of percent total on the gradient.

### Fluorescence Microscopic Analysis of GFP-G1 Distribution

Cells expressing GFP-G1 were plated on coverslips in 35 mm dishes or 6-well plates at 1 × 10^6^ cells per dish. At 48h, the samples were washed in PBS, fixed 45 min in 3% paraformaldehyde in 0.1M sodium phosphate, quenched in 50 mM glycine in PBS, blocked and permeabilized in 2.5% goat serum + 0.1% Triton-X100 in PBS (or 2.5% goat serum + 0.1% saponin in PBS) and then incubated with 1° (1h room temperature) and 2° (30 min room temperature) antibodies diluted in the blocking medium with intervening washes in 0.1% Triton-X100 (or saponin) in PBS. The antibodies indicated in the figures served as reference markers. After staining, the coverslips were washed in 0.1% Triton-X100 (or saponin) in PBS and finally in PBS before mounting in Prolong Gold Antifade (Invitrogen/ThermoFisher). Coverslips were imaged using a Nikon C1 laser scanning confocal unit attached to a Nikon Eclipse TE2000-E microscope with a 100X, 1.45-numerical-aperature (NA) Plan Apochromat objective. Image analysis for Tf recycling was performed using NIS-Elements Software (Nikon) to manually define regions of interest on confocal images. All other image analyses, Tf uptake and colocalization experiments, were preformed using Image J with JACoP plug-in. For live cell imaging, the cells were plated very similarly but in optical dishes (MatTek) and were examined after changing from culture medium to Krebs-Ringer-Bicarbonate-Hepes medium.

### Incubations of cells at 20 °C and with inhibitors of protein synthesis and degradation

To test the effects of decreased temperature on the distribution of GFP-G1, cells plated on coverslips were incubated at 20 °C in a temperature-controlled incubator under CO_2_ for 2 or 4h and were fixed and processed for fluorescence microscopy alongside a control maintained at 37 °C. Following fixation, the samples were immunostained for reference markers of the TGN (Golgin 97, proinsulin) and ERC (transferrin receptor).

To track ABCG1 turnover following inhibition of protein synthesis, INS1 cells were plated 1 × 10^6^ cells per dish in 35 mm dishes and then used at 48h. Cycloheximide 20 μg/ml was added at 0 time and inhibitors of proteasomal degradation (10 μM MG132) or lysosomal degradation (200 nM bafilomycin A) or both together were added to selected sets of plates. At specified timepoints, the plates were washed twice in chilled PBS, lysed in 1% Triton, 10 mM Tris, 100 mM NaCl, 1 mM EDTA, pH 7.4 and proteinase inhibitor cocktail, cleared of insoluble material by centrifugation and prepared for protein assay (BCA) and western blotting. Calpeptin 1 μM was also tested as an inhibitor of turnover in separate experiments.

### Pulse-Chase Biosynthetic Labeling and Immunoadsorption

INS1 cells were preincubated 30 min in cysteine-and methionine-free RPMI medium containing 1.0 mg/ml BSA, pulse labeled 30 min in the same medium containing ^35^S Easy Tag Express Protein Labeling Mix (Perkin Elmer NEG772014; 120 μCi/ml) and then washed and chase incubated in low glucose (5.5 mM) DMEM containing 2 mM glutamine and 1% FBS. Individual wells were used for each timepoint and media were removed, spun in a microcentrifuge to clear any cellular debris. Cells were washed twice with PBS, lysed 20 min on ice with lysis buffer (1% Triton X-100, 10 mM Tris, 100 mM NaCl, 1 mM EDTA and proteinase inhibitors), and cleared of debris by centrifugation. An aliquot of each supernatant was used to measure protein and another aliquot was diluted with RIPA buffer and preadsorbed with Protein A-Sepharose 30 min at 4 °C and then used for immunoadsorption overnight at 4 °C with rabbit anti-ABCG1. The amount of anti-ABCG1 (2 μl) sufficient to adsorb all radiolabeled transporter was determined by initial titration. Following washing three times with 1% Triton containing lysis buffer and once with PBS, immunoadsorbed label was solubilized in sample buffer and resolved by SDS-PAGE on 12% gels. After fixation in isopropanol-acetic acid, the gels were dried and used for phosphorimaging and quantification using the Fujifilm MultiGauge program.

### RNAi-mediated knockdown of ABCG1 in INS1 cells and analysis of uptake and trafficking of fluorescent transferrin and fluorescent cholera toxin

For RNAi-mediated knockdowns, INS1 cells were transfected using the Lipofectamine RNAiMax transfection reagent (Invitrogen/ThermoFisher) and a reverse transfection procedure. Complexes of 50 pmol siRNA/lipofectamine were generated in Optimem and incubated 30 min at room temperature in 35 mm dishes. Cells suspended in antibiotic-free growth medium (1×10^6^ per dish) were added to complexes and cultured 48 h. At 48 h, medium was changed to standard growth medium; at 72 h cells were used for experiments. siRNAs were Dharmacon ON-TARGETplus.

- Control (Ctl) ON-TARGETplus Non-targeting Pool (D-001810-10)
- ABCG1 ON-TARGETplus Rat ABCG1 SMARTpool (GeneID 85264) (Cat# L-093864-02)
- ABCG1 3’UTR ‑ ON-TARGETplus Custom Duplex

Sense AGGCAAAACCGGAGAAGAAUU
Antisense 5’-P-UUCUUCUCCGGUUUUGCCUUU

Assessment of the extent of knockdown was performed by western blotting and was found to be ~90% loss of endogenous ABCG1 (Figure S1 in [27]).

To examine endocytic uptake and recycling, Control-and ABCG1-knockdown INS1 cells were seeded during knockdown on coverslips and used between 72–96h following start of knockdown. For uptake experiments, the cells were starved 30 min at 37 °C in serum-free DMEM containing 0.1% BSA, chilled on an ice-water slurry and then labeled 15 min at 0 °C with 10 μg/ml rat transferrin-Dylight488 and 20 nM cholera toxin-Alexa568. Samples were then washed with the same medium without label and warmed to 37 °C in fresh medium. At specified timepoints, the samples were washed in PBS, fixed 45 min in 3% paraformaldehyde in 0.1M sodium phosphate and processed for immunstaining and observation by fluorescence microscopy as described above under Fluorescence Microscopic Analysis of GFP-G1 Distribution.

For transferrin recycling experiments, cells that had been knocked down and starved as above were incubated 30 min at 37 °C with 10 μg/ml rat transferrin-Dylight488 in DMEM-0.1% BSA. Following washing, the medium was replaced with DMEM-BSA containing 0.25 mg/ml unlabeled transferrin. At specified timepoints, the cells were fixed 30 min in paraformaldehyde as above, then washed 3x in HBSS buffer, cell surface-labeled 10 min with 5 μg/ml wheat germ agglutinin-Alexa555 in HBSS, washed further in HBSS, then PBS and mounted, observed and analyzed as above.

### Procedure for ELISA Assay

Cell-associated insulin (extracted in 1% Triton, 10 mM Tris, 100 mM NaCl, proteinase inhibitors with 1 min bath sonication) and secreted insulin were assayed by ELISA using a homemade kit. An anti-rat proinsulin/insulin Mab (Meridian Life Sci E86201M) was used to coat the plates; anti-rat insulin antibody (Millipore 1013K) was used as sandwich antibody; and insulin standards were Millipore 8013-K.

### Western Blotting

For western blotting, samples were electrophoresed on NuPAGE 4–12% Bis-Tris Gels (Invitrogen/ThermoFisher) and transferred to nitrocellulose. Blotting with primary and secondary antibodies was done in 2.5% milk in PBS-0.1% Tween20. Secondary antibodies conjugated to IR (infrared; Licor) were used for detection; dilutions for Licor secondary reagents were: 1:10,000 for goat anti-rabbit 696 and donkey anti-goat 800 and 1:25,000 for goat anti-mouse 800. Quantitative densitometry of saved Tiff images was done using Fujifilm MultiGauge software. For nearly all blots of clarified cell lysates (as in turnover studies), γ-adaptin was also probed and used for normalization among the different samples. The exception was when blotting for TfR (mobility nearly the same as γ-adaptin) where actin was probed for normalization. For sucrose gradient fractions, the samples analyzed were the same portion of the total of each fraction enabling calculation of fractional distribution on the gradient.

### Statistical Analyses

Data were analyzed using statistical software in GraphPad Prism 7 for unpaired Student’s t-tests. In all figures, * p<0.05; *** p<0.001, **** p<0.0001. All results with error bars are presented as mean ± SEM.

## Acknowledgments

We are grateful to Dr. James Casanova for advice and discussions and for critically reading the manuscript. The authors appreciate the highly valued earlier contributions of Drs. Jeffrey Sturek (University of Virginia School of Medicine) and Lynn Hedrick (La Jolla Institute of Allergy and Immunology) in exploring the role of ABCG1 in insulin secretion. The earlier misdiagnosis of ABCG1 localization was made in the Castle lab and JDC accepts full responsibility.

## Supplementary Material

**Fig S1.** Uncut western blots for Fig 1.

**Fig S2.** Full x-ray film for immunoadsorption of ^35^S-labeled ABCG1 shown in Fig 4A. Full scatter plot for the colocalization data presented in Fig 4E.

**Fig S3.** Full western blots showing results from three (out of four) independent experiments where level of TfR was compared in Control and ABCG1 knockdown samples.

**Fig S4.** (A) Example images of Tf-DyLight488 and CTxB-Alexa555 and immunostained GM130 in Control and ABCG1-depleted cells at 5, 15, and 30 min; (B) example images of immunostained a-mannosidase-II with either TfR or EEA1.

**Video 1.** Colocalization and co-migration of LC3-mRFP and GFP-G1. 35 frames were shot at 1 frame/2 sec. Movie plays at 5 fps.

**Video 2.** Robust trafficking of GFP-G1 toward and away from the cell periphery. 40 frames were shot at 1 fps. Movie plays at 5 fps.

**Additional Supporting Information.** All data for seven turnover experiments for Fig 4B grouped by treatment; Scatter plots for Fig 5C-E.

